# Diversity of Sericostomatidae (Trichoptera) Caddisflies in the Montseny Mountain Range

**DOI:** 10.1101/057844

**Authors:** Jan Niklas Macher, Martina Weiss, Arne J. Beermann, Florian Leese

**Affiliations:** Aquatic Ecosystem Research, University of Duisburg-Essen; Department of Animal Ecology, Evolution and Biodiversity, Ruhr University Bochum

**Keywords:** Population structure, cryptic species, caddisflies, genetic diversity, freshwater invertebrates

## Abstract

Biodiversity is under threat by the ongoing global change and especially freshwater ecosystems are under threat by intensified land use, water abstraction and other anthropogenic stressors. In order to monitor the impacts that stressors have on freshwater biodiversity, it is important to know the current state of ecosystems and species living in them. This is often hampered by lacking knowledge on species and genetic diversity due to the fact that many taxa are complexes of several morphologically cryptic species. Lacking knowledge on species identity and ecology can lead to wrong biodiversity and stream quality assessments and molecular tools can greatly help resolving this problem. Here, we studied larvae of the caddisfly family Sericostomatidae in the Montseny mountains on the Iberian Peninsula. We expected to find cryptic species and that species would not occur in syntopy due to different ecological niches. We sampled 44 stream sites and sequenced 247 larval specimens for the barcoding region of the mitochondrial cytochrome c oxidase gene. A modeling approach was used to assess the bioclimatic preferences of the found species. Two molecular groups were identified. One could be assigned to *Schizopelex furcifera* and one to *Sericostoma* spp. We did find both taxa in syntopy in >50% of sampling sites and could show that the taxa prefer similar bioclimatic conditions. A reexamination of larval specimens showed that *Sericostoma* and *Schizopelex* larvae could not be unambiguously identified to the genus level. Overall, our results show the importance of including molecular tools into biodiversity assessments in order to correctly identify the species diversity of a region and to prevent wrong assessment results.

## Introduction

Biodiversity is under threat by multiple stressors such as climate change and intensified anthropogenic land use (Vörösmarty et al. 2010, Steffen et al. 2015). To understand how stressors impact on biodiversity, it is crucial to inventorise the current state of ecosystems as well as the number and genetic diversity of species living in them. This is especially important for freshwater ecosystems, as they provide humanity with highly valuable drinking water and food resources. Worldwide, there are ongoing efforts to assess the state of freshwaters by monitoring their biodiversity (e.g. in the EU, see Kallis & Butler 2001). These efforts are however hampered by the fact that many freshwater taxa are complexes of several so-called cryptic species (e.g. Weiss et al. 2014, Obertegger et al. 2014, Katouzian et al. 2016) and are difficult to identify (Haase et al. 2006). With the rise of molecular tools during the last decade, it is now straightforward to identify species by sequencing the barcoding region of the cytochrome c oxidase one gene (Hebert et al. 2003). While the knowledge of actual species numbers and how to identify these species is valuable in it own right, it is also important to understand these species’ ecologies and their response to environmental change if they are to be effectively used in biomonitoring programs. This has been rarely done for cryptic species complexes (but see e.g. Pfenninger et al. 2003, Rissler et al. 2007, Macher et al. 2016a, Gabaldon et al. 2016 for examples). Knowledge on species’ ecologies is urgently needed to improve monitoring programs and understand how species and ecosystems are impacted by the ongoing environmental change.

Here, we studied larvae of the common caddisfly family Sericostomatidae (Trichoptera) in the Montseny Mountain Range on the Iberian peninsula (Figure 1a). The Montseny is located at the intersection of the arid and warm climate of the mediterranean lowlands to the west and the cool, precipitation-rich climate of the mountainous region to the east (Thuiller et al. 2003). Catchments within the Montseny are characterized by steep altitudinal gradients, with altitude increasing from below 300 masl to 1706 masl within approximately 10 kilometres and therefore, climatic variables are highly variable as well ( see Jump et al. 2007, Penuelas & Boada 2003). Since caddisflies are merolimnic organisms and can actively disperse between habitats via their winged adult stages (e.g. Geismar et al. 2014), their distribution within a small area is expected to be defined by ecological demands. On the Iberian Peninsula, species of the caddisfly genera *Sericostoma* and *Schizopelex* occur in sympatry (González & Martínez 2011, Ruiz-Garcia et al. 2014). *Sericostoma* is known to consists of an yet unknown number of morphologically cryptic species (Malicky 2005). While adults of *Sericostoma* and *Schizopelex* can be readily distinguished, the larvae of some species within these genera are notoriously difficult or impossible to distinguish from each other based on morphology (Malicky 2005, Vieira-Lanero 2000, Waringer & Graf 2013). They therefore form a cryptic larval complex, a phenomenon also known from other taxa (Pfenninger et al. 2007, Webb & Suter 2011, Carew et al. 2007, Jackson et al. 2014, Zhou et al. 2007). Due to the highly variable environmental conditions in the Montseny, the area seemed highly suitable to study small scale diversity and occurrence of sympatric species.

**Figure 1:**
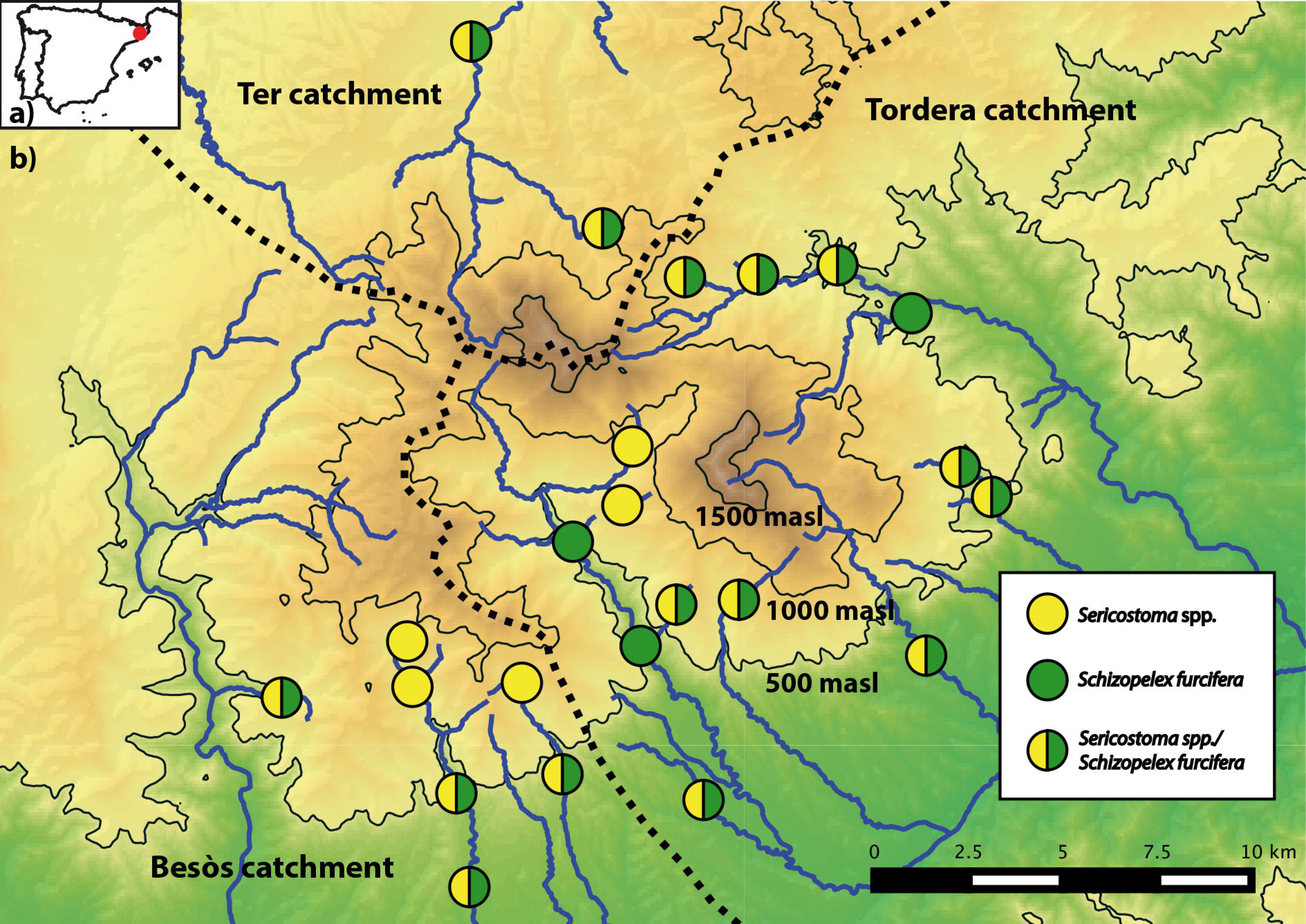
a) Position of the Montseny mountains on the Iberian Peninsula indicated by a red dot. b) Sampling sites in the Montseny and presence of Sericostoma spp., Schizopelex furcifera or both taxa.

We expected 1) to find morphologically cryptic species sericostomatid larvae in the Montseny mountain range, 2) that different Sericostomatidae species seldom co-occur in syntopy due to different ecological demands and 3) that species show a population structure corresponding to altitude, but not between catchments due to their good dispersal ability.

To test these hypotheses we analysed the mitochondrial cytochrome c oxidase gene subunit 1 (CO1) to determine the number and distribution of Sericostomatidae species found in the Montseny. In a second step, we used a modeling approach based on bioclimatic variables to find possible explanations for the occurrence of species and third, we performed population genetic analyses to investigate gene flow between populations and thus infer the dispersal abilities of the different species.

## Materials and Methods

### Sampling and data

Samples from 44 sites were taken during a field trip to the Montseny in September 2013, covering the three main watersheds of the Montseny (Tordera, Besòs, Ter) and an altitudinal gradient from 120 masl to 1295 masl (Figure 1b). Caddisfly larvae were collected using kick nets (HydroBios, Kiel). All larvae were stored in 70% ethanol in the field, transferred to 96% ethanol after less than six hours and stored at 4°C until further analysis. Larvae of Sericostomatidae were identified as belonging to the *Sericostoma/Schizopelex* larval complex using the key in Waringer & Graf (2013) under a stereo microscope. Sericostomatidae larvae were found in 24 sites (see Table A1). QGIS (v 2.8) was used to create distribution maps. A sampling permit for protected areas (Parc Natural del Montseny) was obtained from the park management prior to sampling.

### DNA extraction, amplification, and sequencing

For Sericostomatidae larvae, DNA was extracted from the body wall of 247 larval specimens (1 – 15 specimens per site, 24 sampling sites) using a salt extraction protocol (Sunnucks and Hales 1996). A 841-bp fragment of the barcoding gene COI was amplified using the primers HCO_mod and LCO_mod (Leese et al. 2005, modified after Folmer et al. 1994). PCR mix was prepared using the following protocol: 1 × PCR buffer, 0.2 mM dNTPs, 1 μl of DNA template, 0.025 U/μl Hotmaster Taq (5 PRIME GmbH, Hilden, Germany), 0.5 μM of each primer. The mix was filled up to 25 μl with sterile H_2_O and placed in a thermocycler for amplification. PCR settings for the COI amplification were: initial denaturation at 94°C for 2 min; 36 cycles of denaturation at 94°C for 20 s, annealing at 46°C for 30 s, extension at 65°C for 60 s; final extension at 65°C for 5 min. 9 μl of the PCR product were purified enzymatically with 10 U of Exonuclease I and 1 U Shrimp Alkaline Phosphatase (Thermo Fisher Scientific, Waltham) by incubating at 37°C for 25 min and a denaturation step at 80°C for 15 min. Bidirectional sequencing was performed on an ABI 3730 sequencer by GATC Biotech (Constance, Germany).

### Species delimitation

Geneious 6.0.5. (Biomatters) was used to assemble raw reads and the MAFFT plugin (v. 7.017 Katoh and Standley, 2013) was used to compute a multiple sequence alignment (automatic algorithm selection, default settings). The final length of the cropped alignments was 841 bp for Sericostomatidae. The alignment was translated into amino acids using translation table 5 (invertebrate mitochondrial codon usage table) to make sure that no stop codons were present. The best model of evolution for further analyses of the data was selected with jModeltest 2.1.2 (Darriba et al., 2012) (default settings). Fabox (Villesen 2007) was used to collapse sequences into haplotypes. PopART (v.1, popart.otago.ac.nz) was used to create statistical parsimony haplotype networks (Clement et al. 2000) with a 95% connection limit. To determine the number of species in the *Sericostoma/Schizopelex* larval complex, both the tree-based Generalized Mixed Yule Coalescent (GMYC) approach (Pons et al. 2006) and the automated distance-based barcode gap determination (ABGD, Puillandre et al., 2012) approaches were used. For GMYC analyses, an ultrametric tree for all unique COI haplotypes was calculated using BEAST v.1.8.0 (Drummond et al. 2012). BEAST was run for 10 million MCMC generations, sampling every 100th tree and using both standard coalescent and the GTR + G sequence evolution model. Tracer v.1.6 (Rambaut et al. 2013) was used to test for effective sampling size (ESS) and convergence of parameters. TreeAnnotator v.1.8 (Rambaut and Drummond 2013) was used to generate a linearized consensus tree, discarding the first 3000 trees as burn-in. R v. 3.1.1 (R Core Team 2014) was used for analysis of the resulting tree with ‘SPLITS’ (Species Limit by Threshold Statistics) (Ezard et al. 2009) with the single threshold model to test for the presence of multiple species within the dataset. For ABGD analyses, the K2P-model of sequence correction (Kimura 1980). Sequences of all specimens from each molecularly identified group were blasted against the Barcode of Life database (Ratnasingham & Hebert 2007) to verify species assignment. Alignments were created with Geneious and networks were computed with popArt as described above.

### Bioclimatic variables analyses

The bioclimatic preferences of species were modelled using MaxEnt 3.3.3e (Phillips et al. 2004) and for the same geographic area as in Macher et al. (2016b).

## Results

### Molecular species delimitation

A total of 248 Sericostomatidae larvae from 24 sites were barcoded for the COI barcoding gene. The 841 bp long COI alignment for the Sericostomatidae larvae had 137 (16.3%) variable sites and a GC content of 35%. The null model of a single species was rejected both with the GMYC approach (likelihood ratio for single threshold model: 31.68, p < 0.001) and the ABGD approach (Pmax 0.77-3.59%). ABGD suggested the presence of two groups, but GMYC found four groups in the caddisfly dataset. Blast search assigned 134 specimens to *Schizopelex furcifera* (ABGD and GMYC group 1) and 113 specimens (ABGD group 2, GMYC groups 2, 3, 4) to different *Sericostoma* species (*Sericostoma personatum*, *S. flavicorne*, *S. vittatum*, *S. pyrenaicum*, *S. galeatum*) with no clear assignment of species names to molecularly identified groups. The three subgroups of *Sericostoma* that were found by GMYC as being separate species were thus further treated as *Sericostoma* spp., since Leese et al. (unpublished) could show that COI cannot be used for reliable species delimitation in the genus *Sericostoma*. Afterwards, the species found by the GMYC and ABGD approaches are referred to as *Schizopelex furcifera* and *Sericostoma* spp. 30 larvae each of *Sericostoma* spp. and *Schizopelex furcifera* were again checked with the identification key by Vieira-Lanero (2000) to see whether an identification was possible with the knowledge about clade affiliation of the specimens. However, this was not the case as all features mentioned in the key varied within and between species.

### Bioclimatic characterization

As in Macher et al. (2016b), the 6 variable model resulted in good AUC values for both *Schizopelex furcifera* and *Sericostoma* spp. (0.82 and 0.84, respectively), thus it was chosen for further analyses to mediate between lower variable correlation and high model fitting. *Schizopelex furcifera* occurrence was best predicted by the variables Temperature Annual Range and Precipitation of Coldest Quarter, as was the occurrence of *Sericostoma* spp. Bioclimatic niche overlap was 0.876 for *Schizopelex furcifera* and *Sericostoma* spp. The range overlap computed for occurrence likelihoods of >50% was 0.826 for *Schizopelex furcifera* and *Sericostoma* spp.

## Discussion

In this study, we investigated the distribution and bioclimatic niches of Sericostomatidae caddisfly species in the Montseny mountain range. We expected to find morphologically indistinguishable species in larvae of the trichoptera family Sericostomatidae in the Montseny mountain range. Indeed, at least two species of Sericostomatidae were found to occur in the Montseny. In the *Sericostoma/Schizopelex* larval complex, at least two molecular clades were found, one of which could be molecularly assigned to *Schizopelex furcifera*, a species that is endemic to the North-Western Iberian Peninsula and whose larvae have not yet been described (Waringer & Graf 2013.). The other clade belongs to the *Sericostoma* species complex, which is known to consist of several undescribed species with unknown distribution ranges (Malicky 2005). We could not assign the found *Sericostoma* larvae to a species name due to highly inaccurate results when comparing our molecular data with that deposited in the Barcode of Life database by other researchers. It is thus evident that the genus *Sericostoma* needs to be revised using a combination of morphological (of both larvae and imago) and molecular data, a task that is beyond the aim of this study. Our second expectation was that different Sericostomatidae species seldom co-occur in syntopy due to different ecological demands. This expectation was not met for the sericostomatid caddisflies. *Sericostoma spp*. and *Schizopelex furcifera* were often found in syntopy with no apparent pattern regarding altitudinal preferences. The genus *Sericostoma*, however, is known to inhabit a wide range of stream and river habitats in all of Europe, often occurring in the upper and middle reaches of streams (Leese et al. 2005).

According to the MaxEnt models, the factors explaining the occurrence of both *Sericostoma* spp. and *Schizopelex furcifera* are precipitation of the colder season and the temperature range throughout the year. For aquatic species, higher precipitation means that more water is available throughout the year, which is especially important in generally dry areas or areas with high precipitation seasonality. It seems thus possible that *Sericostoma* spp. and *Schizopelex furcifera* are relying on constant flow of the streams they inhabit. We stress the point that all data presented here is based on a very limited number of specimens only and needs to be interpreted with care, also due to the fact that nothing is known about the frequency and extent of dispersal events in Sericostomatidae species. Studies involving nuclear markers and possibly a greater number of specimens per site are needed to verify the observed patterns of genetic variation on a geographically small scale and also to clarify the taxonomic status of *Sericostoma* spp. Our results show the potential of molecular techniques to study species diversity and ecology of species. Molecular studies are highly valuable to understand the impact of environmental change and stressors on biodiversity (Pauls et al. 2013, Balint et al. 2011). As many bioassessment and monitoring programs worldwide rely on species occurrence data and species’ ecological traits as a metric to measure ecosystem quality (e.g. Carter & Resh 2001, Haase et al. 2004, Stark 2001), molecular tools can help to speed up and improve accuracy of species identification. This is urgently needed to make sure that biomonitoring programs use the right information and that the right conclusions are drawn from the data generated. In combination with data that can inform on ecology of species (e.g. remote sensing data and experiments as in Elbrecht et al. 2016), molecular data can greatly improve assessment of stream health and biomonitoring programs. This is important, since potentially costly restoration measures are based on biomonitoring data and a lot of money might be lost in case of wrong biomonitoring results. In addition, only by using molecular methods the different genetic strains of species in different habitats be found and effectively protected, which helps choosing the most valuable populations and target conservation efforts.

## Acknowledgments

We thank the Parque Natural Montseny for sampling permissions, Nuria Bonada for help with organising permissions and discussions and Alexander Weigand for helpful discussions.

## Data accessibility

CO1 DNA sequences:

GenBank accession numbers for *Sericostoma* sp.: KX365492 - KX365604

GenBank accession numbers for *Schizopelex furcifera.:* KX365605 - KX365738

## References

Bálint, M., Domisch, S., Engelhardt, C., Haase, P., Lehrian, S., Sauer, J., Theissinger, K., Pauls, S. and Nowak, C., 2011. Cryptic biodiversity loss linked to global climate change. Nat. Clim Change 1, 313–318.

Carew, M. E., Pettigrove, V., Cox, R. L., & Hoffmann, A. A. (2007). DNA identification of urban Tanytarsini chironomids (Diptera: Chironomidae). Journal of the North American Benthological Society, 26(4), 587–600.

Carter, J. and Resh, V., 2001. After site selection and before data analysis: sampling, sorting, and laboratory procedures used in stream benthic macroinvertebrate monitoring programs by USA state agencies. J N Am Benthol Soc 20, 658–682.

Clement, M., Posada, D. and Crandall, K., 2000. TCS: a computer program to estimate gene genealogies. Mol Ecol 9, 1657–1659.

Darriba, D., Taboada, G., Doallo, R. and Posada, D., 2012. jModelTest 2: more models, new heuristics and parallel computing. Nat Methods 9, 772.

Drummond, A., Suchard, M., Xie, D. and Rambaut, A., 2012. Bayesian phylogenetics with BEAUti and the BEAST 1.7. Mol Biol Evol 29, 1969–1973.

Elbrecht, V., Beermann, A. J., Goessler, G., Neumann, J., Tollrian, R., Wagner, R., … Leese, F. (2016). Multiple⁏stressor effects on stream invertebrates: a mesocosm experiment manipulating nutrients, fine sediment and flow velocity. Freshwater Biology.

Ezard, T., Fujisawa, T. and Barraclough, T., 2009. SPLITS: species’ limits by threshold statistics. R Package Version 1.

Haase, P., Lohse, S., Pauls, S., Schindehütte, K., Sundermann, A., Rolauffs, P. and Hering, D., 2004. Assessing streams in Germany with benthic invertebrates: development of a practical standardised protocol for macroinvertebrate sampling and sorting. Limnol.-Ecol. Manag. Inland Waters 34, 349–365.

Haase, P., Murray-Bligh, J., Lohse, S., Pauls, S., Sundermann, A., Gunn, R., & Clarke, R. (2006). Assessing the impact of errors in sorting and identifying macroinvertebrate samples. In The Ecological Status of European Rivers: Evaluation and Intercalibration of Assessment Methods (pp. 505–521). Springer Netherlands.

Hebert, P. D., Cywinska, A., & Ball, S. L. (2003). Biological identifications through DNA barcodes. Proceedings of the Royal Society of London B: Biological Sciences, 270(1512), 313–321.

Folmer, O., Black, M., Hoeh, W., Lutz, R. and Vrijenhoek, R., 1994. DNA primers for amplification of mitochondrial cytochrome c oxidase subunit I from diverse metazoan invertebrates. Mol Mar Biol Biotechnol 3, 294–299.

Gabaldón, C., Fontaneto, D., Carmona, M. J., Montero-Pau, J., & Serra, M. (2016). Ecological differentiation in cryptic rotifer species: what we can learn from the Brachionus plicatilis complex. Hydrobiologia, 1–12.

Geismar, J., Haase, P., Nowak, C., Sauer, J., & Pauls, S. U. (2015).Local population genetic structure of the montane caddisfly Drusus discolor is driven by overland dispersal and spatial scaling. Freshwater Biology, 60(1), 209–221.

González, M. A., & Martínez, J. (2011). Checklist of the caddisflies of the Iberian Peninsula and Balearic Islands (Trichoptera). Zoosymposia, 5, 115–135.

Jackson, J. K., Battle, J. M., White, B. P., Pilgrim, E. M., Stein, E. D., Miller, P. E., & Sweeney, B. W. (2014). Cryptic biodiversity in streams: a comparison of macroinvertebrate communities based on morphological and DNA barcode identifications. Freshwater Science, 33(1), 312–324.

Jump, A., Hunt, J. and Penuelas, J., 2007. Climate relationships of growth and establishment across the altitudinal range of Fagus sylvatica in the Montseny Mountains, northeast Spain. Ecoscience 14, 507–518.

Kallis, G., & Butler, D. (2001). The EU water framework directive: measures and implications. Water policy, 3(2), 125–142.

Katoh, K. and Standley, D., 2013. MAFFT multiple sequence alignment software version 7: improvements in performance and usability. Mol Biol Evol 30, 772–780.

Katouzian, A. R., Sari, A., Macher, J. N., Weiss, M., Saboori, A., Leese, F., & Weigand, A. M. (2016). Drastic underestimation of amphipod biodiversity in the endangered Irano-Anatolian and Caucasus biodiversity hotspots. Scientific reports, 6.

Kimura, M., 1980. A simple method for estimating evolutionary rates of base substitutions through comparative studies of nucleotide sequences. J Mol Evol 16, 111–120.

Leese, F., & Wagner, R. (2005). The" Sericostoma-problem"-molecular, chemotaxonomic, and autecological approaches (Trichoptera: Sericostomatidae). Lauterbornia, 54, 161–163.

Macher, J. N., Salis, R. K., Blakemore, K. S., Tollrian, R., Matthaei, C. D., & Leese, F. (2016a). Multiple-stressor effects on stream invertebrates: DNA barcoding reveals contrasting responses of cryptic mayfly species. Ecological Indicators, 61, 159–169.

Macher, J. N., Weiss, M., Beermann, A., & Leese, F. (2016b). Cryptic diversity and population structure at small scales: The freshwater snail Ancylus (Planorbidae, Pulmonata) in the Montseny mountain range. bioRxiv, doi: 054551.

Malicky, H. 2005. Ein kommentiertes Verzeichnis der Köcherfliegen (Trichoptera) Europas und des Mediterrangebiets. Linzer Biologische Beiträge, 37, 533–596

Múrria, C., Morante, M., Rieradevall, A., Ribera, A. and Prat, N, 2014. Genetic diversity and species richness patterns in Baetidae (Ephemeroptera) in the Montseny Mountain range (North-East Iberian Peninsula). Limnetica 33, 313–326.

Obertegger, U., Flaim, G., & Fontaneto, D. (2014). Cryptic diversity within the rotifer Polyarthra dolichoptera along an altitudinal gradient. Freshwater Biology, 59(11), 2413–2427.

Pauls, S., Theissinger, K., Ujvarosi, L., Balint, M. and Haase, P., 2009. Patterns of population structure in two closely related, partially sympatric caddisflies in Eastern Europe: historic introgression, limited dispersal, and cryptic diversity 1. J N Am Benthol Soc 28, 517–536.

Peñuelas, J. and Boada, M., 2003. A global change⁏Linduced biome shift in the Montseny mountains (NE Spain). Glob. Change Biol. 9, 131–140.

Pfenninger, M., Staubach, S., Albrecht, C., Streit, B. and Schwenk, K., 2003. Ecological and morphological differentiation among cryptic evolutionary lineages in freshwater limpets of the nominal form⁏group Ancylus fluviatilis (O.F. Müller, 1774). Mol Ecol 12, 2731–2745.

Pfenninger, M., & Schwenk, K. (2007). Cryptic animal species are homogeneously distributed among taxa and biogeographical regions. BMC evolutionary biology, 7(1), 1.

Phillips, S. J., Dudík, M., & Schapire, R. E. (2004, July). A maximum entropy approach to species distribution modeling. In Proceedings of the twenty-first international conference on Machine learning (p. 83). ACM.

Phillips, S. J., & Dudík, M. (2008). Modeling of species distributions with Maxent: new extensions and a comprehensive evaluation. Ecography, 31(2), 161–175.

Pons, J., Barraclough, T., Gomez-Zurita, J., Cardoso, A., Duran, D., Hazell, S., Kamoun, S., Sumlin, W. and Vogler, A., 2006. Sequence-based species delimitation for the DNA taxonomy of undescribed insects. Syst Biol 55, 595–609.

Puillandre, N., Lambert, A., Brouillet, S. and Achaz, G., 2012. ABGD, Automatic Barcode Gap Discovery for primary species delimitation. Mol Ecol 21, 1864–1877.

Rambaut, A. and Drummond, A., 2013. TreeAnnotator v1. 7.0.

Rambaut, A., Suchard, M., Xie, D. and Drummond, A., 2013. Tracer v1.5, Available from beast.bio.ed.ac.uk/Tracer.

Ratnasingham, S. and Hebert, P., 2007. BOLD: the Barcode of Life Data System (www.barcodinglife.org). Mol Ecol Notes 7, 355–364.

Rissler, L. and Apodaca, J., 2007. Adding more ecology into species delimitation: ecological niche models and phylogeography help define cryptic species in the black salamander (Aneides flavipunctatus). Syst. Biol. 56, 924–942.

Ruiz-Garcia, A., & Ferreras-Romero, M. (2014). A new species of genus Schizopelex McLachlan (Trichoptera, Sericostomatidae), from the southern Iberian Peninsula. Zootaxa, 3866(2), 297–300.

Stark, J. D., 2001. Protocols for sampling macroinvertebrates in wadeable streams. Cawthron Institute, New Zealand

Steffen, W., Richardson, K., Rockström, J., Cornell, S., Fetzer, I., Bennett, E., Biggs, R., Carpenter, S., de Vries, W. and de Wit, C., 2015. Planetary boundaries: Guiding human development on a changing planet. Science 347, 1259855.

Sunnucks, P. and Hales, D., 1996. Numerous transposed sequences of mitochondrial cytochrome oxidase I-II in aphids of the genus Sitobion (Hemiptera: Aphididae). Mol Biol Evol 13, 510–524.

Thuiller, W., Vayreda, J., Pino, J., Sabate, S., Lavorel, S. and Gracia, C., 2003. Large⁏scale environmental correlates of forest tree distributions in Catalonia (NE Spain). Glob. Ecol. Biogeogr. 12, 313–325.

Vieira-Lanero, R. (2000). Las larvas de los tricopteros de Galicia (Insecta: Trichoptera). Universidad de Santiago de Compostela, Santiago de Compostela.

Villesen, P., 2007. FaBox: an online toolbox for fasta sequences. Mol Ecol Notes 7, 965–968.

Vörösmarty, C., McIntyre, P., Gessner, M., Dudgeon, D., Prusevich, A.,Green, P., Glidden, S., Bunn, S., Sullivan, C., Reidy Liermann, C. and Davies, P., 2010. Global threats to human water security and river biodiversity. Nature 467, 555–561.

Waringer, J. & Graf, W. (2011) Atlas der mitteleuropäischen Köcherfliegenlarven—Atlas of Central European Trichoptera Larvae. Erik Mauch Verlag, Dinkelscherben, Germany, 468 pp

Webb, J. M., & Suter, P. J. (2011). Identification of larvae of Australian Baetidae. Museum Victoria Science Reports, 15, 1–24.

Weiss, M., Macher, J., Seefeldt, M. and Leese, F., 2014. Molecular evidence for further overlooked species within the Gammarus fossarum complex (Crustacea: Amphipoda). Hydrobiologia 721, 165–184.

Zhou, X., Kjer, K. M., & Morse, J. C. (2007). Associating larvae and adults of Chinese Hydropsychidae caddisflies (Insecta: Trichoptera) using DNA sequences. Journal of the North American Benthological Society, 26(4), 719–742.

